# Immunomodulatory mechanisms of submicron phosphatidylserine-exposing polymeric particles (PSPs)

**DOI:** 10.64898/2026.08.07.743390

**Authors:** Maria Thea Rane Dela Cruz Clarin, Kenichi Kimura, Ahmed Nabil, Koichiro Uto, Eri Motoyama, Hnin Htet Htet Aung, Mitsuhiro Ebara, Hiromi Yanagisawa

## Abstract

Macrophages are highly dynamic cells that maintain tissue homeostasis by regulating both initiation and resolution of inflammation. During efferocytosis, macrophages recognize the ‘eat me’ signal, phosphatidylserine (PS), exposed at the surface of apoptotic cells, leading to the resolution of inflammation and acquisition of a pro-resolving phenotype. Inspired by this endogenous mechanism, PS-based biomaterials have demonstrated immunomodulatory potential. However, the molecular mechanisms underlying PS-mediated macrophage reprogramming remain poorly understood. Here, submicron PS-exposing polymeric particles (PSPs; ∼300 nm) were developed to improve the suitability of PSP formulations for future systemic administration while preserving their immunomodulatory activity. PSPs were efficiently internalized by macrophages through both actin- and dynamin-dependent pathways. PSP treatment significantly reduced IL-6 and IL-12p70 production in LPS-stimulated macrophages, whereas induction of the classical anti-inflammatory M2 marker CD206 was limited. Transcriptomic analysis revealed coordinated attenuation of inflammatory signaling pathways, including downregulation of *Myd88, Nfkb1, Rel,* and *Irf8*, together with activation of NRF2-associated antioxidant pathways characterized by increased expression of *Nfe2l2, Hmox1, Prdx1, Gclm, and Gclc.* Activation of antioxidant-associated genes together with reduced *Irf8* expression suggests that PSP promotes inflammatory resolution through coordinated redox adaptation and selective attenuation of inflammatory signaling. Collectively, these findings provide mechanistic insight into PS-mediated macrophage reprogramming and support the future development of systemically administered therapies for chronic inflammatory diseases, including vascular inflammatory disorders.

**Highlights:**

- Submicron PSPs retain immunomodulatory activity of apoptotic cell-mimicking biomaterials.
- PSPs are rapidly internalized through actin- and dynamin-dependent pathways.
- PSPs attenuate inflammatory signaling and selectively suppress IL-6 and IL-12p70 production.
- PSPs induce NRF2-associated antioxidant and glutathione responses.
- Transcriptomics reveals an early redox-adaptive macrophage program.

## 1 Introduction

Inflammation is a fundamental protective response to pathophysiological stimuli that is essential for maintaining tissue homeostasis. Innate immune cells, particularly macrophages, play central roles in orchestrating this response. However, when inflammation becomes chronic, it contributes to the pathogenesis of a wide range of diseases, including cardiovascular diseases such as atherosclerosis and neurodegenerative disorders such as Alzheimer’s and Parkinson’s diseases [1–7]. Inflammation has also emerged as a pathogenic mechanism in hereditary connective tissue disorders. In particular, recent studies have identified macrophage activation and nuclear factor-κB (NF-κB) signaling as early events in Marfan syndrome-associated aortic dissection [8, 9]. Given their role in regulating inflammatory responses, macrophages have emerged as promising therapeutic targets for restoring inflammatory homeostasis.

Among the diverse mechanisms by which macrophages resolve inflammation, efferocytosis is particularly important [10]. This process refers to the clearance of apoptotic cells in a non-inflammatory manner. Apoptotic cells are known to externalize phosphatidylserine (PS) on the cell surface, where it serves as an “eat-me” signal that enables macrophage recognition and engulfment. This promotes the secretion of anti-inflammatory mediators, such as transforming growth factor-β1 (TGF-β1) and interleukin-10 (IL-10), while suppressing the production of pro-inflammatory cytokines IL-6, IL-1β, and tumor necrosis factor-α (TNF-α) [11–13]. These immunomodulatory effects are accompanied by transcriptional reprogramming of macrophages, leading to changes in their functional phenotype and cytokine production.

Inspired by this natural mechanism, apoptotic cell-mimicking biomaterials have been developed to harness the immunomodulatory function of PS such as liposomes, hydrogels, and polymeric systems, for diverse therapeutic applications[14–18]. For instance, PS-functionalized hydrogel with chitosan as a matrix was applied as a wound dressing in a mouse model with diabetic wounds [17]. In another work, the incorporation of PS groups into a methacrylate polymer backbone enabled the preparation of phosphatidylserine-exposing polymeric particles (PSPs) (∼ 1 µm) [19, 20]. These PSPs were developed as an apoptotic cell-mimetic platform for modulating microglial inflammatory responses and demonstrated anti-inflammatory effects following intracerebral administration in an LPS-induced neuroinflammation [19, 20]. While these studies established PSPs as a promising therapeutic platform, the microparticle formulation was designed for local administration and may not be optimal for targeting inflammatory lesions, such as those within the vascular wall, following systemic delivery. Additionally, the mechanisms by which they modulate macrophage function remain poorly understood. These limitations highlight the need to develop a submicron PSP and understand the molecular mechanisms underlying PSP-mediated macrophage modulation, thereby advancing the translational potential of PSPs, particularly for future vascular applications. In particular, whether the PSP induces classical anti-inflammatory macrophage polarization or reprograms macrophages through distinct transcriptional pathways remains unknown. Addressing this knowledge gap is essential for the rational design and optimization of PS-based immunomodulatory biomaterials.

In this study, we developed a submicron PSP formulation and investigated its interaction with macrophages by characterizing cellular uptake, inflammatory responses, and the early transcriptional programs underlying PSP-mediated macrophage modulation.

## 2 Materials and Methods

### 2.1 Materials

2,2’-Azobis(isobutyronitrile) (AIBN), butyl methacrylate (BMA), hydroxyethyl methacrylate (HEMA), *N*, *N*-dimethylformamide (DMF), dichloromethane (DCM), 2-propanol, and imidazole hydrochloride were all purchased from FUJIFILM Wako Pure Chemical Corporation (Osaka, Japan). 4-Cyano-4-[[(dodecylthio)carbonothioyl]thio]pentanoic acid (CDSPA) and trifluoroacetic acid (TFA) were purchased from Tokyo Chemical Industry (Tokyo, Japan). *Tert*-butyl tetraisopropylphosphorodiamidite was obtained from Sigma (St. Louis, MO, USA). *N-*Boc-*L*-serine *tert*-butyl ester was purchased from Watanabe Chemical Industries (Hiroshima, Japan).

### 2.2 Synthesis and characterization of polymer

The poly(BMA-*st*-HEMA) copolymer was synthesized by reversible addition–fragmentation chain transfer (RAFT) polymerization with slight modifications to a previously reported method (**Scheme S1A**) [19]. Briefly, BMA (28.1 mmol), HEMA (7.68 mmol), AIBN (0.04 mmol), CDSPA (0.2 mmol), and DMF (35 mL) were combined in a Schlenk flask. After degassing by nitrogen bubbling for 30 min at room temperature (RT), the reaction mixture was polymerized at 60 °C for 21 h under continuous stirring. The resulting solution was transferred into a dialysis membrane (MWCO 1,000 Da) and dialyzed sequentially against 2-propanol (four changes) followed by DCM (two changes) at RT. Residual solvent was subsequently removed using a rotary evaporator under reduced pressure and further dried under vacuum to obtain the poly(BMA-*st*-HEMA) copolymer.

Poly(BMA-*st*-HEMA-*st*-MPS) was then synthesized via a post-polymerization phosphoramidite reaction (**Scheme S1B**). *Tert-*butyl tetraisopropylphosphorodiamidite (5.0 mmol), *N*-Boc-*L*-serine *tert*-butyl ester (5.0 mmol), imidazole hydrochloride (1.1 mmol), and super-dehydrated DCM (200 mL) were added to a flask and reacted under a nitrogen atmosphere at RT for 21 h. Poly(BMA-s*t*-HEMA) (5.5 mmol hydroxyl groups) was then added to the reaction mixture. Imidazole hydrochloride (14 mmol) was added three times at 45 min intervals, followed by a further 150 min reaction time after the final addition. The reaction mixture was transferred to a dialysis membrane (MWCO 1,000 Da) and dialyzed sequentially against 2-propanol (four solvent exchanges) and DCM (two solvent exchanges) at RT. Solvent removal by rotary evaporation followed by vacuum drying yielded poly(BMA-*st*-HEMA-*st*-(t-Bu/Boc)MPS) as a transparent film. For deprotection, the polymer was dissolved in DCM and treated with 20% TFA. The deprotected polymer was dialyzed against 0.01 M NaOH (two solvent exchanges) followed by deionized water (two solvent exchanges), and subsequently freeze-dried to obtain poly (BMA-*st*-HEMA-*st*-MPS).

The chemical structure of the polymer was confirmed by ^1^H NMR spectroscopy at 400 MHz (JEOL, Tokyo, Japan) using deuterated chloroform as a solvent (**Fig. S1A**).

### 2.3 Preparation of PSP

PS-exposing particles (PSP) composed of poly(BMA-*st*-HEMA-*st*-MPS) were prepared by dialysis-induced self-assembly. The polymer was dissolved in DMF at a concentration of either 1 or 2 mg/mL, with a total solution volume of 20–25 mL. The polymer solution was transferred into regenerated cellulose (RC) dialysis tubing (MWCO 3.5 kDa) and dialyzed against 1 L of deionized (DI) water. The external DI water was replaced at least five times. The resulting PSP suspension was collected and stored at 4 °C until use.

For fluorescent labeling, Nile red (1 wt% relative to the polymer; Fujifilm Wako Pure Chemical Corporation) was dissolved together with the polymer in DMF prior to dialysis. Nile red-loaded PSP were prepared using the same dialysis procedure described above.

Particle suspensions were aliquoted for subsequent characterization, including particle size and zeta potential measurements, nanoparticle tracking analysis (NTA), and cell culture experiments.

### 2.4 Characterization of physicochemical properties

The hydrodynamic diameter and zeta potential of the particles were measured using a dynamic light scattering (DLS) and zeta potential analyzer (ELSZ-2000SZ, Otsuka Electronics, Osaka, Japan). For particle size measurements, 1 mL of the particle suspension was transferred to a disposable cuvette, whereas zeta potential was measured using a flow cell. Particle number was determined using Nanosight NS300 (Nanosight Ltd., Malvern, United Kingdom).

Particle morphology and elemental composition were characterized by scanning transmission electron microscopy coupled with energy-dispersive X-ray spectroscopy (STEM-EDS). Briefly, the particle suspension was deposited onto copper microgrids (HRC-C15, 100 µm pitch; Stem Co., Ltd. and Ouken Shoji Co., Ltd., Japan) and examined using a JEM-2100F microscope (JEOL Ltd., Tokyo, Japan) operated at an accelerating voltage of 200 kV with a 1 nm probe.

To confirm the presence of PS, Annexin V-FITC Apoptosis detection kit (Nacalai Test, Inc.) was used according to the manufacturer’s protocol. Briefly, the samples were diluted using 1X Annexin V binding buffer and 100 µL of each was mixed with 5 µL of Annexin V-FITC conjugate. After incubation for 15 min at RT with protection from light, 400 µL of 1X Annexin V binding solution was added, and the samples were analyzed using a flow cytometer (FACS Melody, BD Biosciences).

### 2.5 Isolation of peritoneal macrophages

C57BL/6J (WT) mice were purchased from The Jackson Laboratory and maintained under specific pathogen-free conditions with a 12 h light/12 h dark cycle. 3 to 6-month-old mice were used. All animal experiments were conducted following animal experimentation guidelines approved by the Institutional Animal Experiment Committee of the University of Tsukuba (approved number 25-398).

Thioglycolate-elicited peritoneal macrophages were isolated as previously described with slight modifications [21]. Briefly, mice received an intraperitoneal injection of 2 mL of 4% thioglycolate broth, and peritoneal cells were harvested by lavage with cold phosphate-buffered saline (PBS) after 3 days. Following red blood cell lysis, cells were washed with PBS, counted using a hemocytometer, and resuspended in RPMI 1640 medium (Gibco, Thermo Fisher Scientific) supplemented with 10% fetal bovine serum (FBS) and 1% antibiotic-antimycotic (Gibco, Thermo Fisher Scientific). Cells were seeded in 12-well plates at a density of 4.4 × 10^5^ cells/well and incubated at 37 °C in a humidified atmosphere containing 5% CO_2_. After 1 h, the medium was replaced to remove non-adherent cells, and the adherent macrophages were cultured overnight before subsequent experiments.

### 2.6 Cell culture experiments

Cells were incubated with lipopolysaccharide (LPS, InvivoGen) at a concentration of 100 ng/mL with or without PSP and collected at the specified time points for subsequent experiments. Cells incubated in medium only were used as a control (CTRL), while treatment with IL-4 (BioLegend) at 200 ng/mL was used to induce M2-like macrophages. For control particles (CP), 300 nm carboxylated beads (24051-10, Polymersciences, Inc.) were used.

For the cellular uptake experiment, cells were treated with LPS, together with CP or Nile Red-loaded PSP at a similar particle concentration (2.15 x 10^10^ particles/mL). For inhibitor experiments, pre-treatment was performed using 10 μM Cytochalasin D (C8273-1MG, Sigma Aldrich) or 100 μM Dynasore (sc-202592, SCB Santa Cruz Biotechnology, Inc.) for 30 min. Consequently, the medium was replaced with treatment conditions with or without the presence of the inhibitor and incubated for 2 h. For each sample, images were taken at three different locations using confocal microscope (LSM980, Carl Zeiss) and quantified using ImageJ.

For the quantification of IL-6, IL-12p70, and TNF-α expression level, the collected supernatant was tested using BD^TM^ Cytometric Bead Array kit (552364, BD Biosciences) according to the manufacturer’s protocol.

For cell viability, cells were seeded in a 96-well plate with the respective treatment groups for 24 h and assessed using Cell Counting Kit-8 (347-07621, Dojindo Laboratories Co., Ltd.). The absorbance was measured at 450 nm using a microplate reader. The cell viability was expressed as a percentage relative to the untreated control group.

### 2.7 mRNA extraction and cDNA synthesis

RNA isolation was performed using RNeasy Plus Micro Kit (QIAGEN), followed by cDNA synthesis using iScript^TM^ Reverse Transcription Supermix (Bio-Rad) according to the manufacturer’s protocol. SYBR Green Master Mix (Takara, Japan) was used for the quantitative real-time PCR (RT-PCR). Quantification of gene expression was performed using the 2^−ΔΔCt^ method, normalized against the housekeeping gene, β-actin, and fold change was reported relative to the untreated control. Corresponding RT-PCR primers are reflected in **Table S1**.

### 2.8 Western Blot

The cell lysates were prepared in RIPA lysis buffer (Sigma-Aldrich) with 1% protease inhibitor (Sigma-Aldrich) and 1% phosphatase inhibitor (Fujifilm Wako Pure Chemical Corporation). Consequently, it was mixed with 3X SDS sample buffer with 2-mercaptoethanol, boiled at 95 °C for 5 min, and subjected to SDS-PAGE. Protein extracts were then transferred to a PVDF membrane (Millipore), blocked, and immunoblotted with primary antibodies in 1:1000 dilution as follows: rabbit anti-GAPDH (2118S, Cell Signaling Technology), goat anti-CD206 (AF2535, R&D Systems). Blocking was performed using either 5% skim milk or 5% Bovine Serum Albumin (BSA) for phosphorylated proteins in 1X TBST. The membrane was incubated for 1 h at RT with anti-rabbit or anti-goat HRP-conjugated secondary antibody (Bio-Rad). Visualization was performed by using chemiluminescence kit (Atto Corporation).

### 2.9 Immunofluorescence (IF) staining

Cells were fixed with 4% paraformaldehyde (PFA) for 5 minutes and washed with 1X PBS three times. Permeabilization was performed using 0.1% Triton X-100 at RT for 10 min, followed by blocking with 5% bovine serum albumin (BSA) for 20 min. The cells were stained with primary antibodies (2 h, RT) at a 1:200 dilution and washed. Cells were stained using the following: mouse anti-CD68 (KP1, Santa Cruz), goat anti-CD206 (AF2535, R&D Systems), rat anti-MHC Class II (M5/114.15.2, Invitrogen), and rat anti-F4/80 (HCA154, Bio-Rad). This was followed by incubation with secondary antibodies conjugated to Alexa Fluor 647 (1:400, Jackson ImmunoResearch) diluted with Hoechst at 1:1000 (Sigma-Aldrich) for 1 h at RT and mounted with Ibidi mounting medium (Ibidi). Images were captured using the confocal microscope (LSM980, Carl Zeiss).

### 2.10 Bulk RNA sequencing (RNA-seq) analysis

WT-derived peritoneal macrophages were incubated for 2 h with LPS or LPS + PSP or without treatment as a control. RNA extraction was performed using Trizol^TM^ Reagent (15596026, Invitrogen) and subsequently sent for sequencing. Libraries were sequenced using the Illumina NovaSeq X Plus platform. The FASTQ files were processed using the CLC genomics Workbench v25.0.3 (QIAGEN) and subsequently used to prepare for the count matrix. Downstream analyses were conducted in R (version 4.4.2) using the DEseq2 (version 1.46.0) package. Differentially expressed genes (DEGs) were determined using an FDR-adjusted p-value < 0.05 and |log fold change| ≥ 0.58.

### 2.11 Statistical Analysis

The GraphPad Prism Version 11.0.0 was used for the preparation of graphs and statistical tests. All data were plotted as mean ± SEM and the Shapiro-Wilk test was used to determine the normality. A two-tailed Student’s t-test or one-way ANOVA was used for datasets that pass the normality, otherwise Mann-Whitney or Kruskal-Wallis test was performed.

## 3 Results

### 3.1 Preparation and physicochemical characterization of PSP

PSPs were prepared from the copolymer by self-assembly using the dialysis method (**Fig. 1A**). To confirm the PS exposure on the PSP surface, the annexin V-FITC binding was evaluated (**Fig. 1B-C** and **Fig. S2A-C**). In comparison with NC (particles without any PS moiety), PSP exhibited a marked rightward shift in the Annexin V-FITC histogram (**Fig. 1B**). This observation is further supported by a significant increase in FITC intensity, confirming the presence of surface-exposed PS following self-assembly (**Fig. 1C**). Surface-exposed PS provides the interface through which PSP are expected to mediate interactions between PSP and macrophages (**Fig. 1D**). STEM-EDS imaging revealed spherical morphology and confirmed the presence of its characteristic elements including carbon, oxygen, phosphorus, and nitrogen, consistent with the expected elemental composition of PSP (**Fig. 1E**). To control the particle size, two different polymer concentrations were used. PSP prepared at 1 mg/mL had a mean diameter of 318 ± 10 nm (**Fig. 1F-G**).

**Fig. 1.**
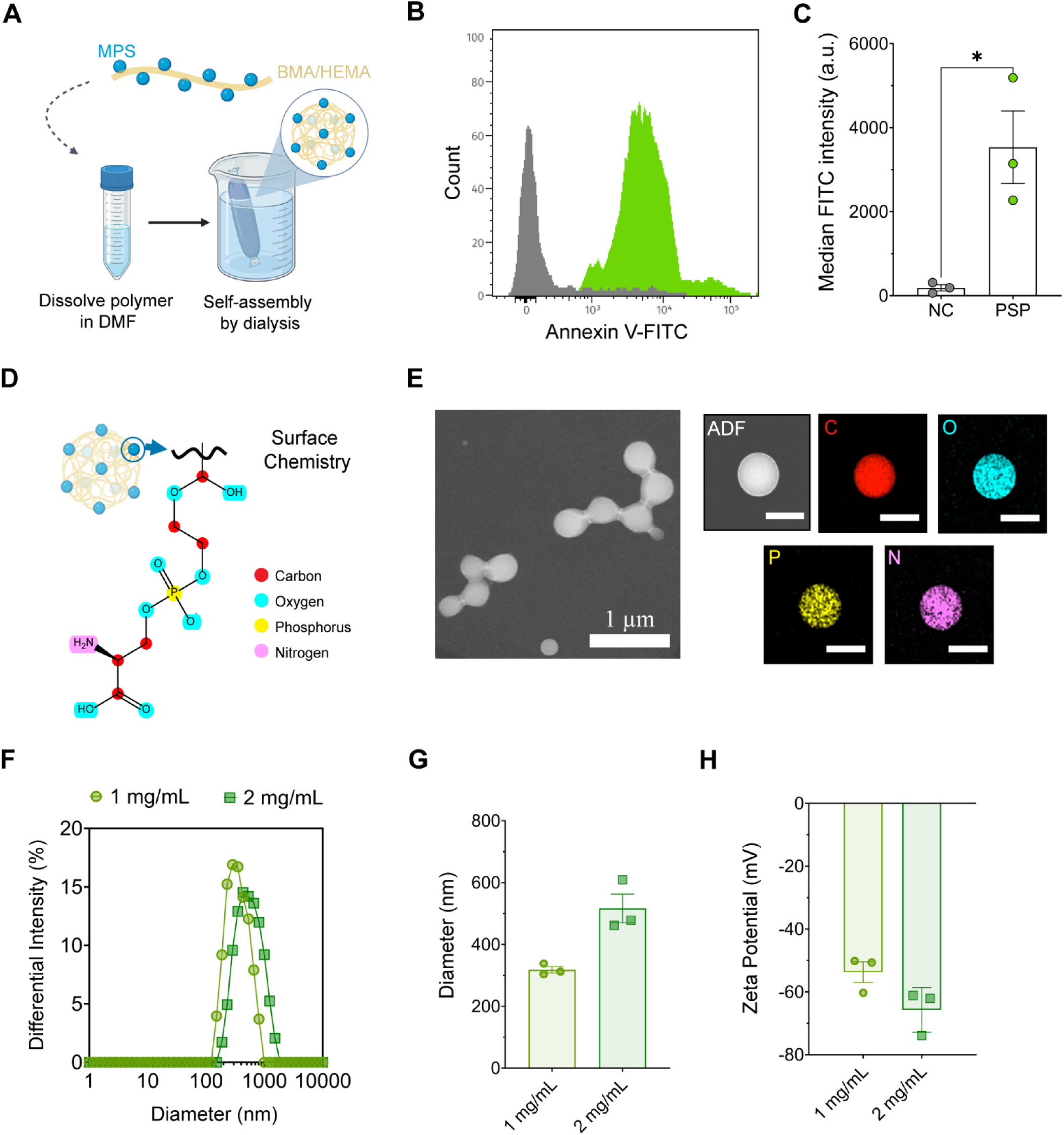
PSP preparation and characterization. **A**, Self-assembly of PSP from poly(BMA-*st*-HEMA-*st*-MPS). **B**-**C,** Annexin V-FITC assay histogram (**B**) and quantified intensity (**C**) to validate PS exposure. Dark grey and lime green represent the negative control (NC) and PSP, respectively. Data represent mean ± SEM (n = 3 independent particle preparations). A two-tailed unpaired t-test. **D**, Surface chemistry of exposed PS. **E**, STEM-EDS imaging of PSP. ADF: Annular dark field. Scale bar = 200 nm. **F-H**, Particle size distribution (**F**), diameter (**G**), and zeta potential (**H**) of PSP formulations prepared at polymer concentrations 1 mg/mL and 2 mg/mL, which yields approximately 300 nm and 500 nm particles. Figure F shows a representative size distribution. Data represent mean ± SEM (n = 3 independent particle preparations).

Compared with the previously reported micron-sized PSP formulation, this represents a substantial reduction in particle diameter [19, 20]. Increasing the polymer concentration to 2 mg/mL, the particle diameter increased to 516 ± 47 nm (**Fig. 1F-G**). Both formulations exhibited a negative surface charge of -54 ± 3 mV and -66 ± 4 mV, respectively (**Fig. 1H**). Collectively, these results demonstrate that PSP can be readily prepared with tunable particle sizes by adjusting polymer concentration while maintaining surface PS exposure and a negatively charged surface. Unless otherwise stated, the 300 nm formulation was used for all subsequent experiments.

### 3.2 PSP selectively suppresses inflammatory cytokine production

To determine whether PSP modulates inflammatory responses in macrophages, thioglycolate-elicited peritoneal macrophages were isolated for subsequent experiments (**Fig. 2A**). The isolated cells predominantly express macrophage markers, F4/80 and CD206, the latter of which is commonly associated with anti-inflammatory macrophages (**Fig. 2B**). In contrast, only a small subset of cells expressed CD68 and MHCII, which are generally associated with pro-inflammatory macrophage. Collectively, these findings are consistent with a homeostatic residential macrophage phenotype (**Fig. 2B**).

**Fig. 2.**
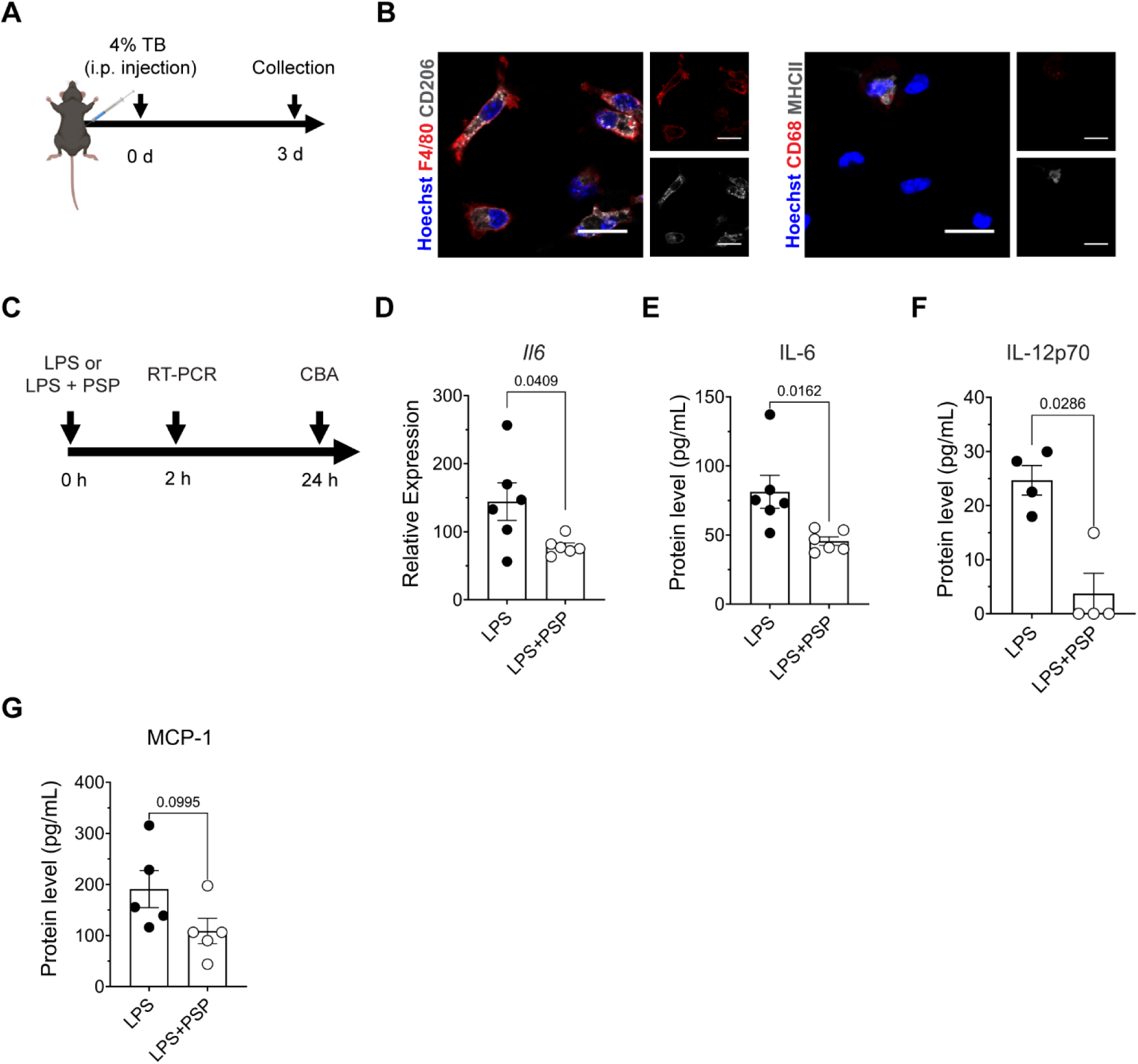
Evaluation of anti-inflammatory activity of PSP in primary peritoneal macrophages. **A**, Isolation of peritoneal macrophages. Mice were intraperitoneally (i.p.) injected with 4% thioglycolate broth (TB) and macrophages were collected after 3 days. **B,** Validation of PM_φ_ identity. Scale bar = 20 µm. **C**, Experimental scheme for the evaluation of PSP. CBA: Cytometric bead array. **D-E**, PSP significantly reduces IL-6 cytokine at mRNA (**D**) and protein (**E**) levels. **F-G**, IL-12p70 (**F**) and MCP-1 (**G**) protein levels. Data represent mean ± SEM (n = 4 to 6 biologically independent animals). All graphs were analyzed using two-tailed unpaired t-test, except for IL-12p70 (Mann-Whitney test).

To determine an optimal working concentration, macrophages were stimulated with LPS in the presence of increasing concentrations of PSP (**Fig. S3A**). PSP reduced *Il6* expression in a concentration-dependent manner, whereas cell viability decreased at the highest concentration tested (53µM) (**Fig. S3B-D**). Based on these results, 38 µM was selected as the working concentration for subsequent experiments (**Fig. 2C**). This concentration is comparable to the effective concentration reported for the previous PSP formulation [20]. PSP significantly reduced *Il6* mRNA expression at 2 h, which was accompanied by a significant reduction in IL-6 protein secretion at 24 h (**Fig. 2D-E**). Likewise, secretion of IL-12p70 was significantly decreased (**Fig. 2F**), whereas MCP-1 showed a decreasing trend (**Fig. 2G**). In contrast, neither *Tnf* nor *Il1b* mRNA expression, nor TNF-secretion was affected by PSP treatment (**Fig. S4A-C**). Together, these findings demonstrate that PSP selectively suppresses inflammatory mediators in LPS-stimulated macrophages.

### 3.3 PSP uptake is mediated by actin- and dynamin-dependent pathways

To characterize the uptake kinetics of PSP, macrophages were incubated with Nile Red-labeled PSP at different time points (**Fig. 3A-C**). At 2 h, approximately 70% of macrophages had internalized PSPs (**Fig. 3A** and **3C**). The intracellular PSP signal continued to increase over time, as shown by the progressive increase in raw integrated density per cell (**Fig. 3A-B**). By 6 h, nearly all macrophages had internalized PSPs, and this level was maintained at 24 h (**Fig. 3C**). Given the rapid uptake of PSPs by macrophages, the uptake mechanism was next investigated using pharmacological inhibitors (**Fig. 3D**). Endocytosis generally involves actin- or dynamin-dependent pathways, which were inhibited by Cytochalasin D and Dynasore, respectively. Treatment with either inhibitor significantly reduced cellular uptake of PSP compared to the control (**Fig. 3E-F**). Collectively, these findings demonstrate that PSPs are rapidly internalized by macrophages through both actin- and dynamin-dependent pathways. Because robust PSP uptake coincided with early suppression of *Il6* expression, subsequent mechanistic analyses were performed at the 2-h time point (**Fig. 2D** and **Fig. 3A-C**).

**Fig. 3.**
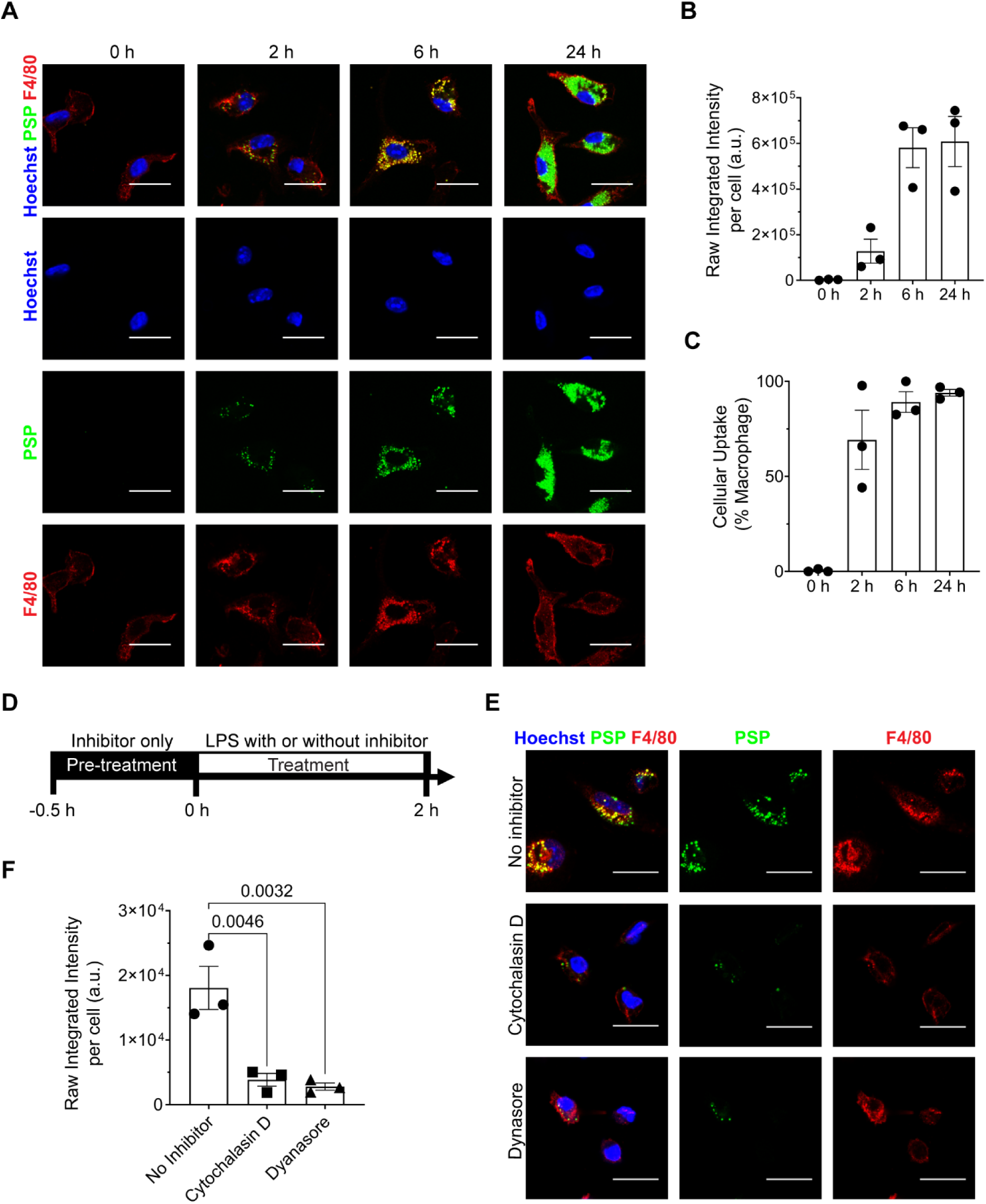
Cellular uptake kinetics and uptake mechanism of PSP in peritoneal macrophages. **A-C,** Macrophage uptake of PSP at different time points: images and quantifications. In Figure 3B, PSP intensity was used for quantification and normalized by cell counts, while Figure 3C shows the percentage of macrophages that uptake PSP. Data represent means ± SEM (n = 3 biologically independent animals). **D,** Evaluation of PSP uptake mechanism using Cytochalasin D and Dynasore to inhibit actin and dynamin, respectively. **E-F,** PSP uptake in LPS-stimulated macrophages. Immunostaining images in the presence or absence of inhibitor (**E**) and quantification (**F**). Data represent means ± SEM (n = 3 biologically independent animals). Scale bar = 20 µm. One-way ANOVA was used for the statistical test.

### 3.4 Control particles lacking PS do not alter *Il6* expression

Given that IL-6 was the inflammatory mediator most significantly suppressed by PSP, *Il6* expression was selected as the primary readout to determine whether this effect was attributable to PS rather than nonspecific particle uptake. Macrophages were therefore treated with control particles (CP) of comparable size but lacking the PS moiety at the same particle number as PSP (**Fig. 4A**). In the presence of CP, it did not alter *Il6* expression compared with LPS-treated group (**Fig. 4B**). Nevertheless, CP was efficiently taken up by macrophages to a similar extent as PSP (**Fig. 4C-E**). These findings demonstrate that particle uptake alone is insufficient to suppress *Il6* expression and identify surface PS as the critical determinant of PSP-mediated immunomodulation.

**Fig. 4.**
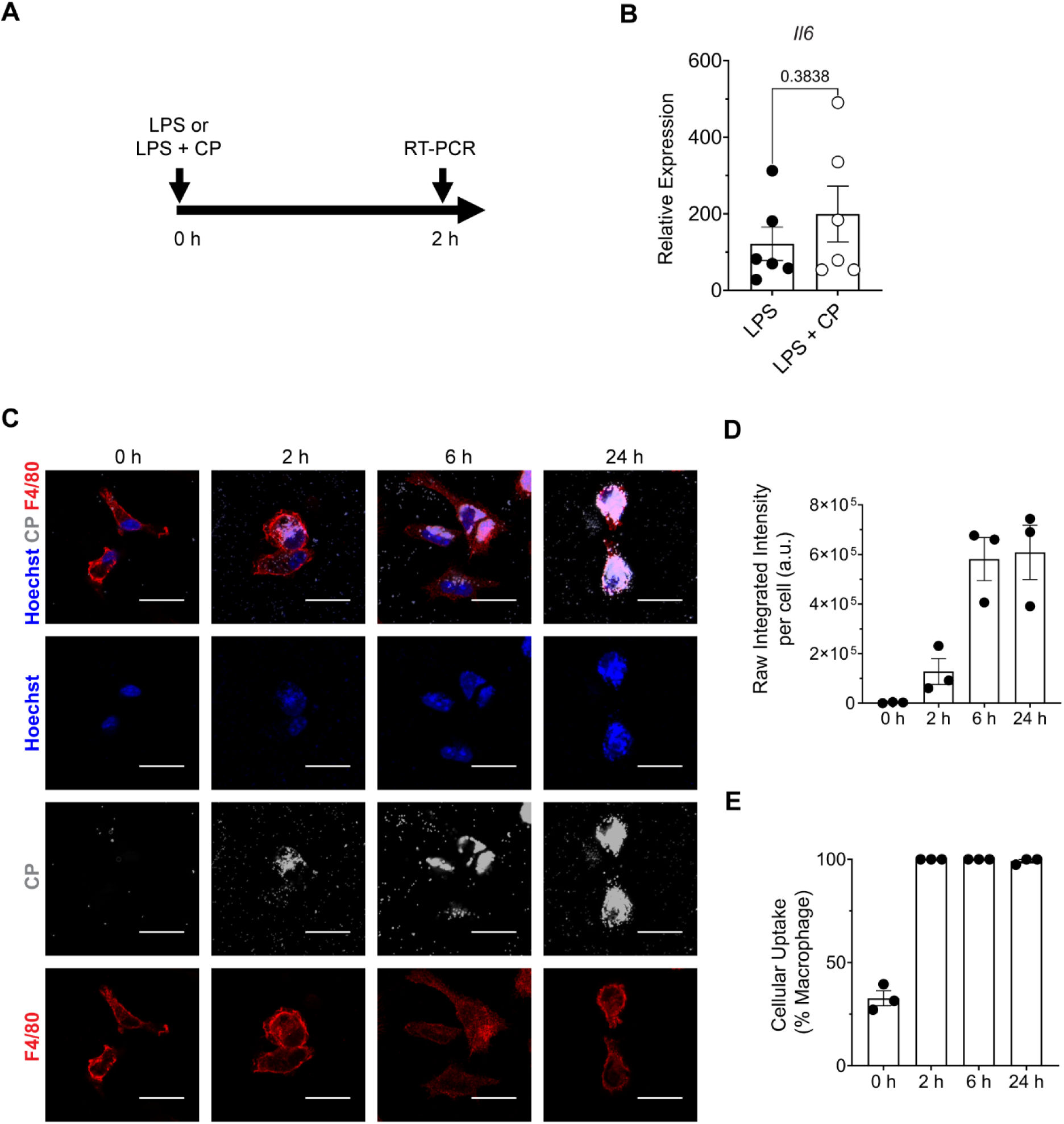
Macrophages uptake control particles without reduction in the *Il6* level. **A**, Evaluation scheme of control particles (CP). **B**, CP treatment did not reduce the *Il6* level in LPS-stimulated macrophages. Data represent means ± SEM (n = 6 biologically independent animals). Mann-Whitney test was used to analyze the graph. **C-E,** Macrophages uptake of CP at different timepoints: images and quantifications. In Figure D, CP intensity was used for quantification and normalized by cell counts, while Figure D shows the percentage of macrophages that uptake CP. Data represent means ± SEM (n = 3 biologically independent animals). Scale bar = 20 µm.

### 3.5 PSP suppresses inflammatory cytokine production with limited induction of the classical M2 marker CD206

Given that suppression of inflammatory cytokines is often associated with classical M2 polarization, CD206 expression was evaluated following PSP treatment (**Fig. S5A**). IL-4, a positive control for M2 polarization, induced robust CD206 expression at all evaluated time points (**Fig. S5B-C**). In contrast, control and LPS-treated macrophages exhibited minimal changes in CD206 expression, whereas a modest increase became detectable at 72 h with PSP (**Fig. S5B-C**). Western blot analysis at 72 h showed that IL-4 markedly increased CD206 protein expression, whereas no significant difference was observed between the LPS and PSP groups (**Fig. S5D-E**). Collectively, these findings indicate that PSP induces only limited expression of the classical M2 marker CD206 under the present experimental conditions and that PSP may regulate macrophage reprogramming through mechanisms distinct from classical M2 polarization.

### 3.6 PSP reprograms macrophages toward an antioxidant-associated transcriptional state

To identify transcriptional programs regulated by PSP, RNA-sequencing (RNA-seq) was performed at the 2 h time point. Principal component analysis (PCA) and unsupervised hierarchical clustering demonstrated a clear separation of LPS-treated and untreated groups, indicating distinct transcriptomic profiles among the groups (**Fig. S6A-B**). Consistently, LPS treatment induced a total of 2,510 differentially expressed genes (DEGs) when compared to the CTRL group (**Fig. S6C**). The upregulated DEGs (1323 genes) correspond to biological pathways such as *innate immune response*, *NF-kB signaling*, *Tnf production*, and *pattern recognition receptor signaling*, indicating the robust induction of inflammatory pathways (**Fig. S6D**). In the presence of PSP, PCA showed a shift in the transcriptomic profile relative to LPS treatment alone (**Fig. S6A)**, and hierarchical clustering of the most variable genes further distinguished LPS and LPS+PSP groups (**Fig. S6E)**. These data suggest that PSP induces transcriptomic changes in macrophages, which may consequently influence their phenotype.

To further investigate the effect of PSP treatment, differential expression analysis between the treated groups (LPS+PSP vs. LPS) identified a total of 220 DEGs (**Fig. 5A**). Pathway enrichment analysis of the DEGs revealed enrichment of antioxidant-related pathways together with reduced inflammatory response pathways in PSP-treated samples (**Fig. 5B-C**). These findings were further substantiated by GSEA, wherein the top-ranked genes showed a high enrichment score in the reactive oxygen species pathway, while a negative enrichment score was observed in a broad set of genes involved in the inflammatory response (**Fig. 5D-E**). At the gene level, PSP treatment increased the expression of oxidative stress-responsive genes, including *Nfe2l2*, *Hmox1*, *Prdx1, Gclc*, and *Gclm* (**Fig. 5F**). In parallel, key inflammatory signaling components such as *Nfkb1* and *Myd88* were reduced, together with marked downregulation of *Stat1*, *Stat2*, and *Irf8* (**Fig. 5G**). Taken together, these transcriptomic analyses demonstrate that PSP not only suppresses inflammatory signaling, but also reprograms macrophages through coordinated activation of the antioxidant pathway and attenuates inflammatory gene networks.

**Fig. 5.**
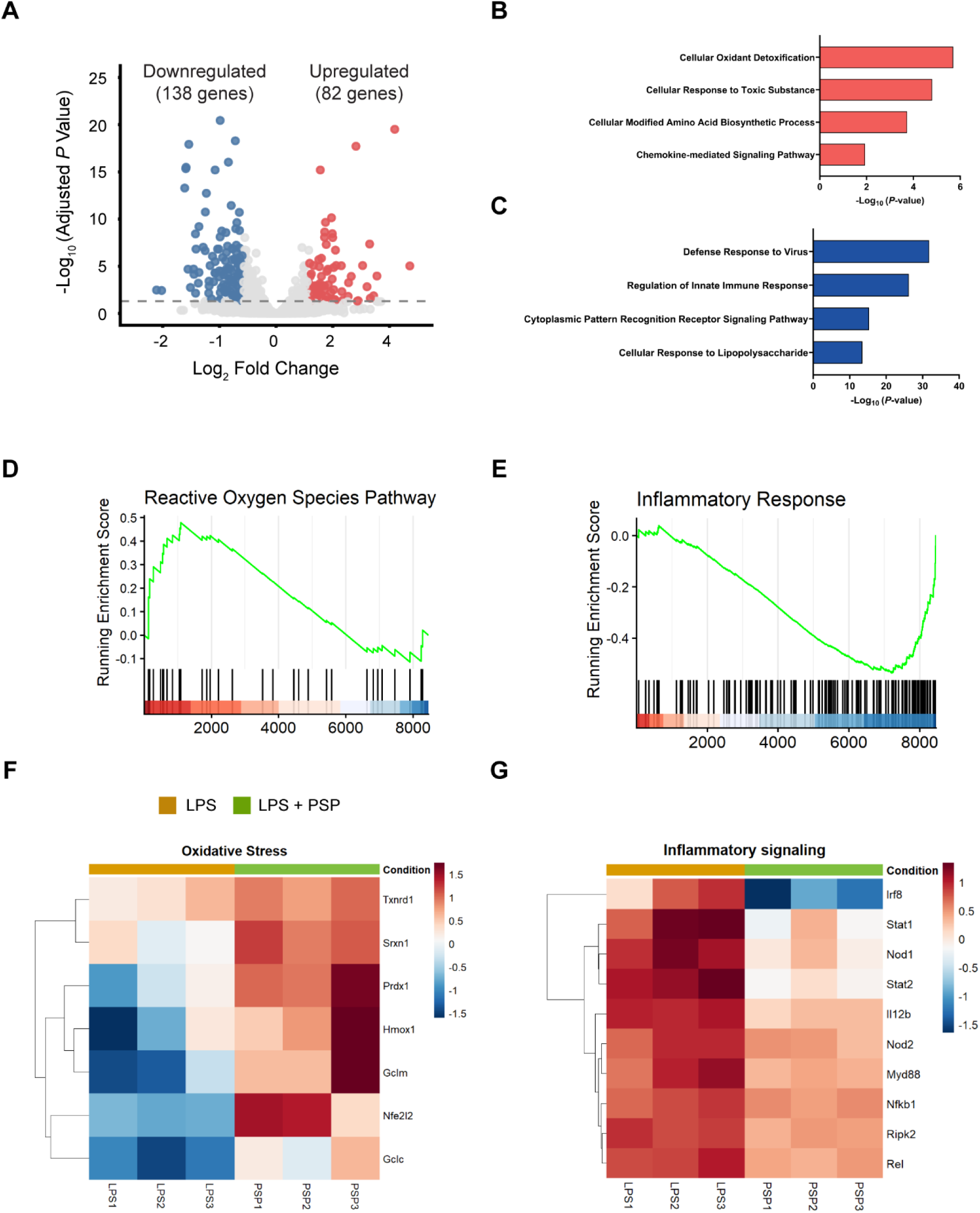
Transcriptomic analysis of PSP- and LPS-treated groups. **A**, Volcano plot showing the DEGs from LPS + PSP vs. LPS. **B-C,** Enriched pathways induced by upregulated (**B**) and downregulated (**C**) DEGs. **D-E**, Gene Set Enrichment Analysis (GSEA) of Reactive Oxygen Species (**D**) and Inflammatory Response (**E**). **F-G**, Heatmap of gene expression related to oxidative stress (**F**) and inflammatory (**G**) gene sets.

## 4 Discussion

In this study, we showed that PSP provides a PS-presenting platform with tunable physical properties that is efficiently internalized by macrophages through actin- and dynamin-dependent pathways to reduce inflammation, while also initiating antioxidant pathways.

Previous studies established PSP as an apoptotic cell-mimicking biomaterial that exhibits anti-inflammatory activity and therapeutic potential following intracerebral administration [19, 20]. Here, the platform was adapted into a submicron formulation to improve its suitability for future systemic applications while maintaining its biological activity. Given the influence of particle size in biodistribution, the control of this parameter is important in material design, particularly for systemic injection where several biological barriers must be overcome to reach the target organ. For instance, nanoparticles with less than 10 nm in diameter are rapidly eliminated through renal and urinary excretion [22, 23]. Moreover, nanoparticles ranging from 80 to 240 nm have been successfully used for imaging the aortic wall [24]. The PSP developed in this study (∼300 nm) represents a rational balance between remaining sufficiently large to avoid rapid clearance while reducing the particle size near the range reported to access the aortic wall. Consequently, this submicron formulation provides a promising platform for the future development of systemically administered therapeutics targeting vascular diseases such as aortic dissection and atherosclerosis. Although in vivo biodistribution was beyond the scope of this study, these findings support the rationale for developing a smaller PSP formulation.

Macrophages are highly plastic immune cells, and their phenotype is dynamically regulated by the cues from their surrounding environment. Consequently, the physicochemical properties of biomaterials, such as surface chemistry, particle size, and cargo can profoundly influence macrophage activation and function [25–27]. In the present study, PSP and CP are efficiently internalized by macrophages, indicating that particle uptake alone is insufficient to explain the observed immunomodulatory activity. Despite comparable physicochemical characteristics and equivalent particle numbers, only PSP reduced *Il6* expression, suggesting that the biological response is highly influenced by the presentation of PS, rather than nonspecific phagocytosis. This observation is consistent with previous findings, in which polymeric particles prepared without the phosphatidylserine-containing monomer (MPS) similarly failed to suppress NF-kB activity [19]. This highlights the importance of PS as a bioactive surface cue for macrophage regulation.

The rapid uptake of PSP by macrophages through both actin- and dynamin-dependent pathways suggests that multiple endocytic mechanisms contribute to the particle internalization. Particle size also influences the internalization pathway. Larger particles (>500 nm) are generally internalized through actin-dependent phagocytic processes, although proteins, including dynamin, may play a role [28, 29]. Given that the present PSP formulation is approximately 300 nm, the involvement of both actin- and dynamin-dependent pathways is biologically reasonable. Similarly, anionic phospholipids and negatively charged PS liposomes have been reported to utilize scavenger receptors such as SR-B1 and GAS6-mediated recognition during uptake [30–32]. Together, these findings suggest that PSP internalization is governed by both its bioactive PS surface and its submicron particle size, allowing rapid macrophage interaction that precedes the observed transcriptional changes.

The RNA-seq analysis revealed that PSP treatment attenuates inflammatory signaling pathways while inducing antioxidant pathways. First, genes involved in toll-like receptor (TLR) signaling and downstream transcriptional programs, including *Myd88*, *Nfkb1*, *Rel*, *Stat1*, *Stat2*, and *Irf8*, were consistently downregulated following PSP treatment. This transcriptomic profile is consistent with the significant reduction in IL-6 and IL-12p70 expression observed in the present study.

IL-12p70 is a pro-inflammatory cytokine composed of two subunits, p35 and p40 (encoded by *Il12a* and *Il12b*, respectively). Previous studies have shown that phagocytosis of apoptotic cells by activated macrophages selectively suppresses *Il12a* and *Il12b* transcription, while exerting comparatively modest effects on other pro-inflammatory cytokines, including *Il1b* and *Il6* [33]. One of the key transcriptional regulators of IL-12p70 is interferon regulatory factor 8 (IRF8), also known as interferon consensus sequence-binding protein (ICSBP). Consistent with this role, ICSBP-deficient mice exhibit markedly impaired IL-12p40 production [34]. In line with this, transcriptomic analysis revealed significant downregulation of *Irf8* in PSP-treated macrophages. Together, these findings suggest that the reduction in IL-12 observed following PSP treatment may be associated with the decreased *Irf8* expression and is consistent with a selective IL-12 regulatory program described during apoptotic cell recognition.

Second, transcriptomic profile induced by PSP treatment showed activation of antioxidant-associated pathways. Among the highly upregulated genes is *Nfe2l2*, which encodes the transcription factor Nrf2 that regulates the cellular antioxidant system through its interactions with antioxidant response elements (ARE) [35]. For instance, it mediates adaptive response against stressors through the increased synthesis rate of glutathione, the endogenous antioxidant in the cells [36]. Moreover, upregulation of NFE2L2 is also reported in tumor-associated macrophages (TAMs) and IL-4-induced M2 macrophages, suggesting its broad role in macrophage reprogramming [37]. Consistent with the activation of this pathway, PSP treatment also increased the expression of several downstream targets of Nrf2, including *Prdx1*, *Hmox1*, *Gclm*, and *Gclc*.

Peroxiredoxin 1 (PRDX1) is a major antioxidant enzyme that detoxifies reactive oxygen species and limits oxidative stress. Recent work demonstrated that LPS-induced macrophage activation is accompanied by reduced acetylation of PRDX1, which potentiates IL-6 production through increased hydrogen peroxide accumulation and ERK signaling [38]. Although PRDX1 acetylation was not examined in the present study, PSP treatment significantly increased *Prdx1* expression, suggesting enhanced antioxidant capacity. Together with the transcriptomic evidence of suppressed inflammatory signaling, these findings raise the possibility that the observed reduction in IL-6 results from both enhanced antioxidant defense and attenuation of inflammatory signaling pathways.

Beyond redox regulation, glutathione metabolism has been implicated in the clearance of apoptotic cells termed efferocytosis [36, 39]. RNAseq analysis showed increased expression of *Glcm*, and *Glcl* with PSP treatment, both of which encode the rate-limiting enzymes for glutathione synthesis and contain ARE regions that can be targeted by Nrf2 [36]. Notably, the loss of the modifier subunit of glutamate cysteine ligase (GCLM) impaired efferocytosis [40]. In addition, *Nrf2*-deficient macrophages aggravate atherosclerosis by promoting inflammation and inhibiting efferocytosis [39]. Together, these findings indicate that PSP may promote a redox-adaptive macrophage state that supports inflammatory resolution.

In light of the limited induction of CD206 observed in this study, these data suggest that macrophage functional reprogramming by PSP may be initiated through metabolic and antioxidant adaptation rather than classical M2 polarization, providing an alternative perspective on the early events that drive inflammatory resolution following macrophage interaction with PSP.

Overall, this study demonstrates that a submicron PS-presenting biomaterial retains the immunomodulatory properties of apoptotic cell while uncovering an early redox-adaptive transcriptional program associated with inflammatory resolution. Functional and transcriptomic analyses revealed coordinated attenuation of inflammatory signaling together with activation of NRF2-associated antioxidant pathways, providing a new perspective that macrophage reprogramming induced by apoptotic cell-mimicking biomaterials may be initiated through metabolic and redox adaptation rather than robust induction of classical M2 markers. These findings advance the mechanistic understanding of phosphatidylserine-based immunomodulatory biomaterials and establish a foundation for future studies evaluating the biodistribution, therapeutic efficacy, and safety of PSP following systemic administration. By harnessing endogenous pathways that promote inflammatory resolution, the mechanistic insights gained in this study provide a foundation for expanding the potential application of PSPs to vascular inflammatory diseases, including Marfan syndrome-associated aortic disease.

## Data Availability

The datasets generated and analyzed during this study are available in the Gene Expression Omnibus (GEO repository) (accession no. GSExxxxxx).

## CRediT authorship contribution statement

**Maria Thea Rane Dela Cruz Clarin:** Methodology, Validation, Formal analysis, Investigation, Data Curation, Visualization, Funding Acquisition, Writing – Original Draft. **Eri Motoyama:** Investigation, Resources. **Hnin Htet Htet Aung:** Investigation, Validation, Formal analysis. **Ahmed Nabil:** Methodology, Supervision. **Koichiro Uto**: Methodology, Supervision. **Kenichi Kimura:** Conceptualization, Methodology, Supervision, Project administration, Funding acquisition, Writing – Review & Editing. **Mitsuhiro Ebara:** Conceptualization, Supervision, Funding acquisition, Project administration, Writing – Review & Editing. **Hiromi Yanagisawa:** Conceptualization, Supervision, Project administration, Funding acquisition, Writing – Review & Editing.

## Acknowledgments

The authors thank the Laboratory Animal Resource Center at the University of Tsukuba for their excellent animal care. The authors also acknowledge the Bioanalysis Unit and Electron Microscopy Unit of the National Institute for Materials Science (NIMS) for providing access to instrumentation, technical support, and electron microscopy analyses. The authors sincerely thank Masanori Toyofuku for providing access to the nanoparticle tracking analysis (NTA) instrument used in this study and Shunsuke Matsumoto for his technical assistance with polymer synthesis. The authors are grateful to Patrick Sips, Julie De Backer, Lynn Y. Sakai, Erna Raja, Keiichi Asano, Marina Matsumiya, and Kosuke Sato for the discussion.

---

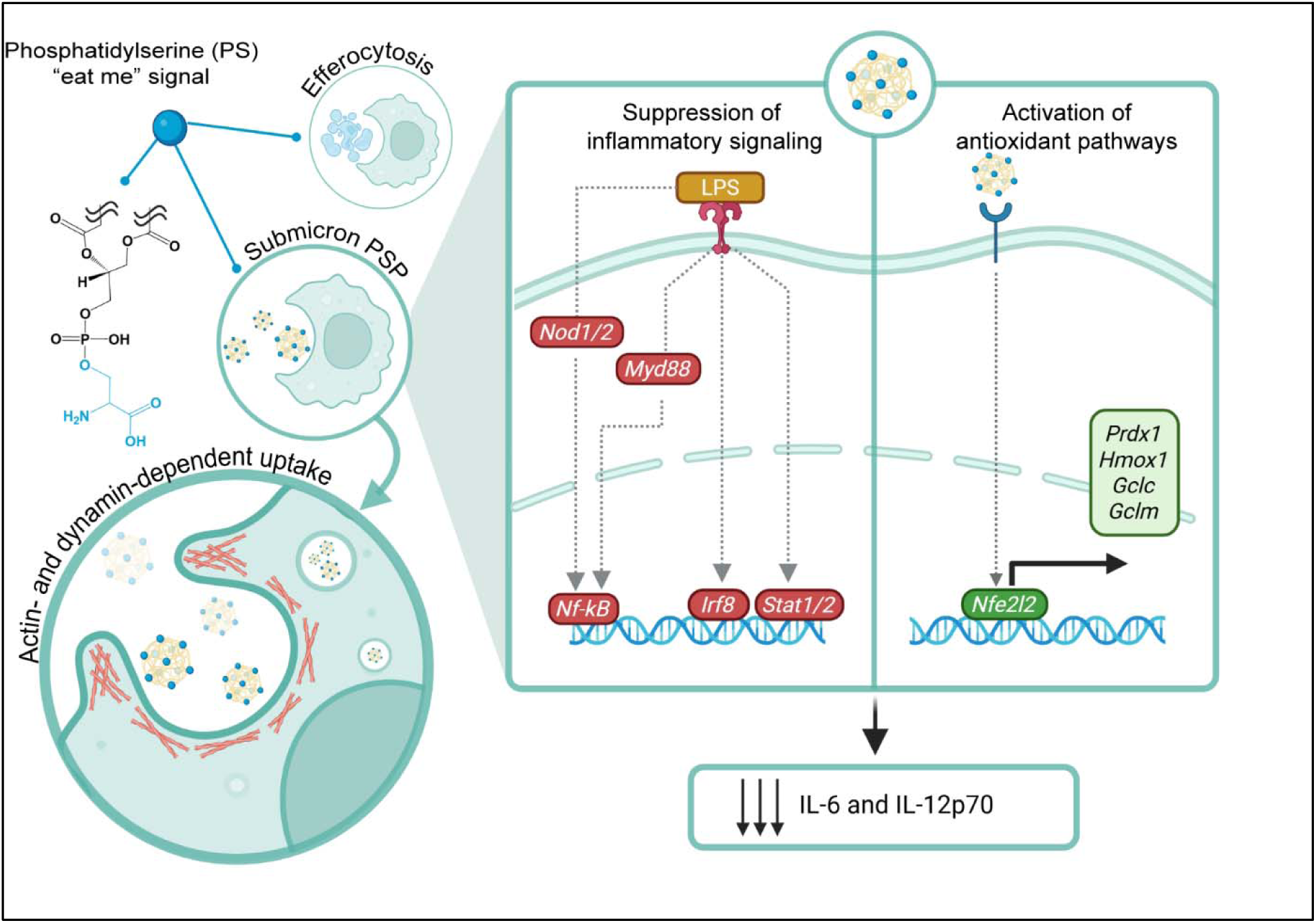

