## Supplementary Figures for "Immunomodulatory mechanisms of submicron phosphatidylserine-exposing polymeric particles (PSPs)"

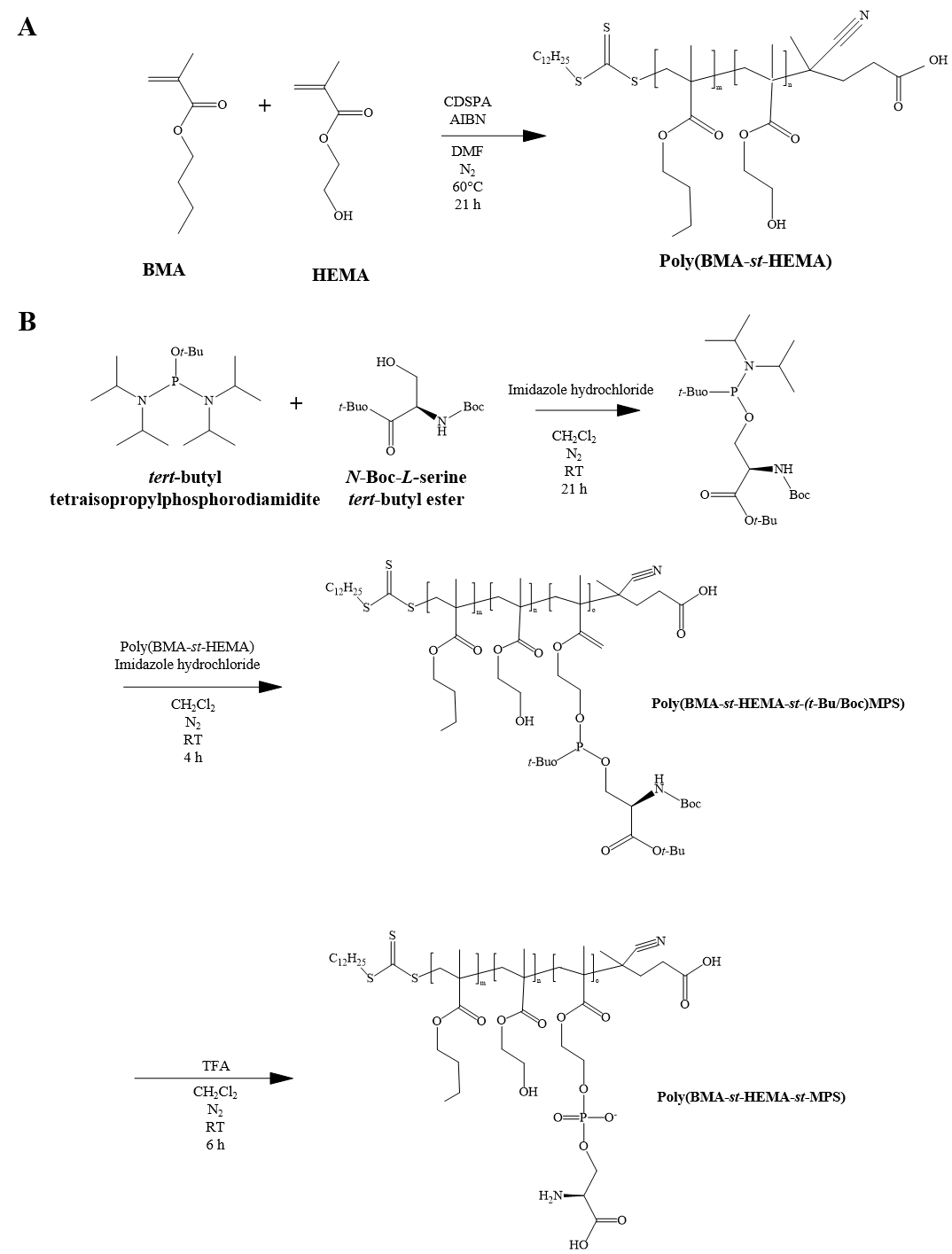

**Scheme S1.** **Synthesis Scheme**. **A,** Synthesis of poly(BMA-*st*-HEMA). **B,** Post-polymerization reaction and deprotection reaction for poly(BMA-*st*-HEMA-*st*-MPS).

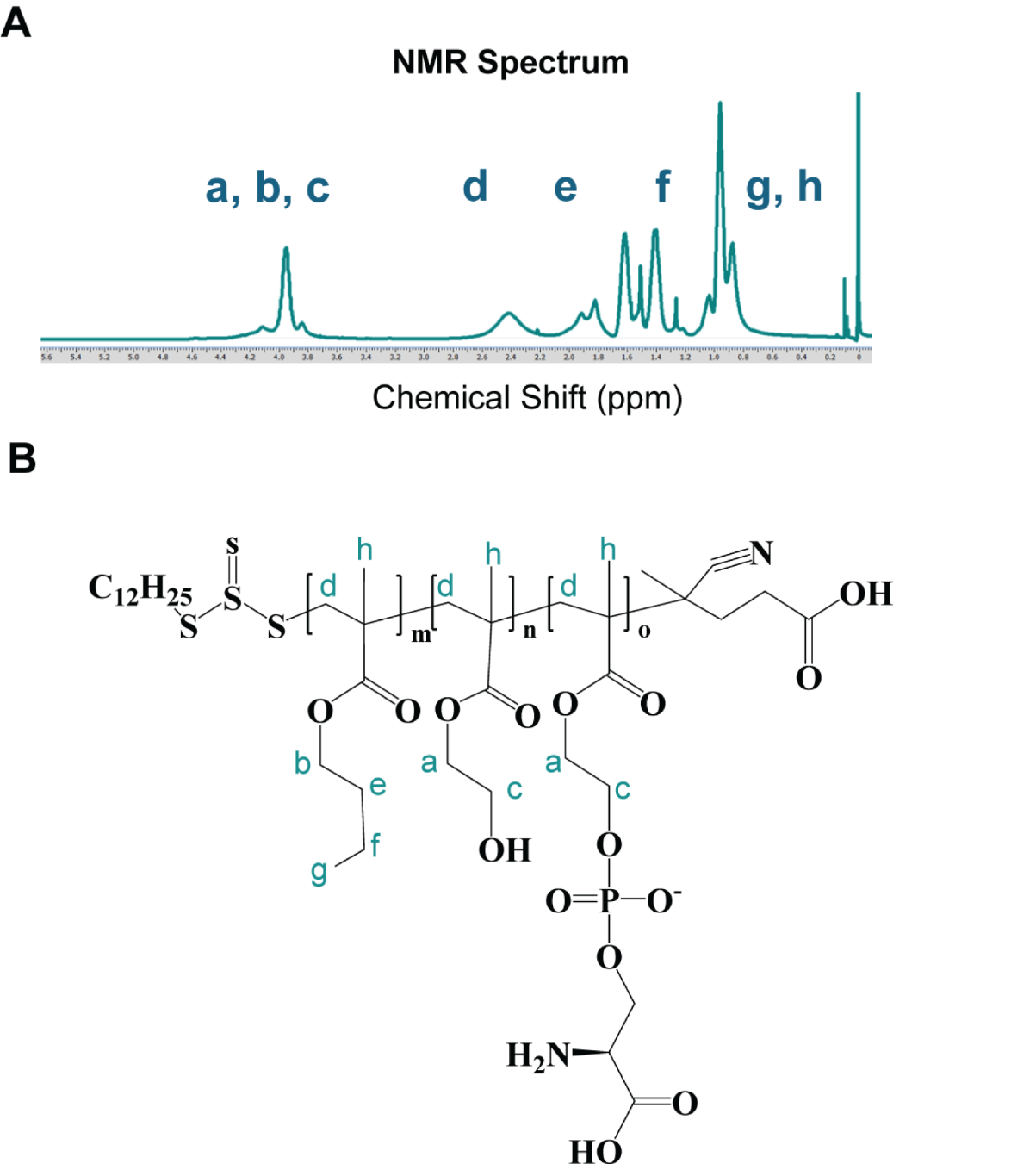

**Fig. S1.** **Polymer characterization**. **A-B**, ^1^H NMR spectrum of poly(BMA-*st*-HEMA-*st*-MPS) (**A**) and its chemical structure (**B**).

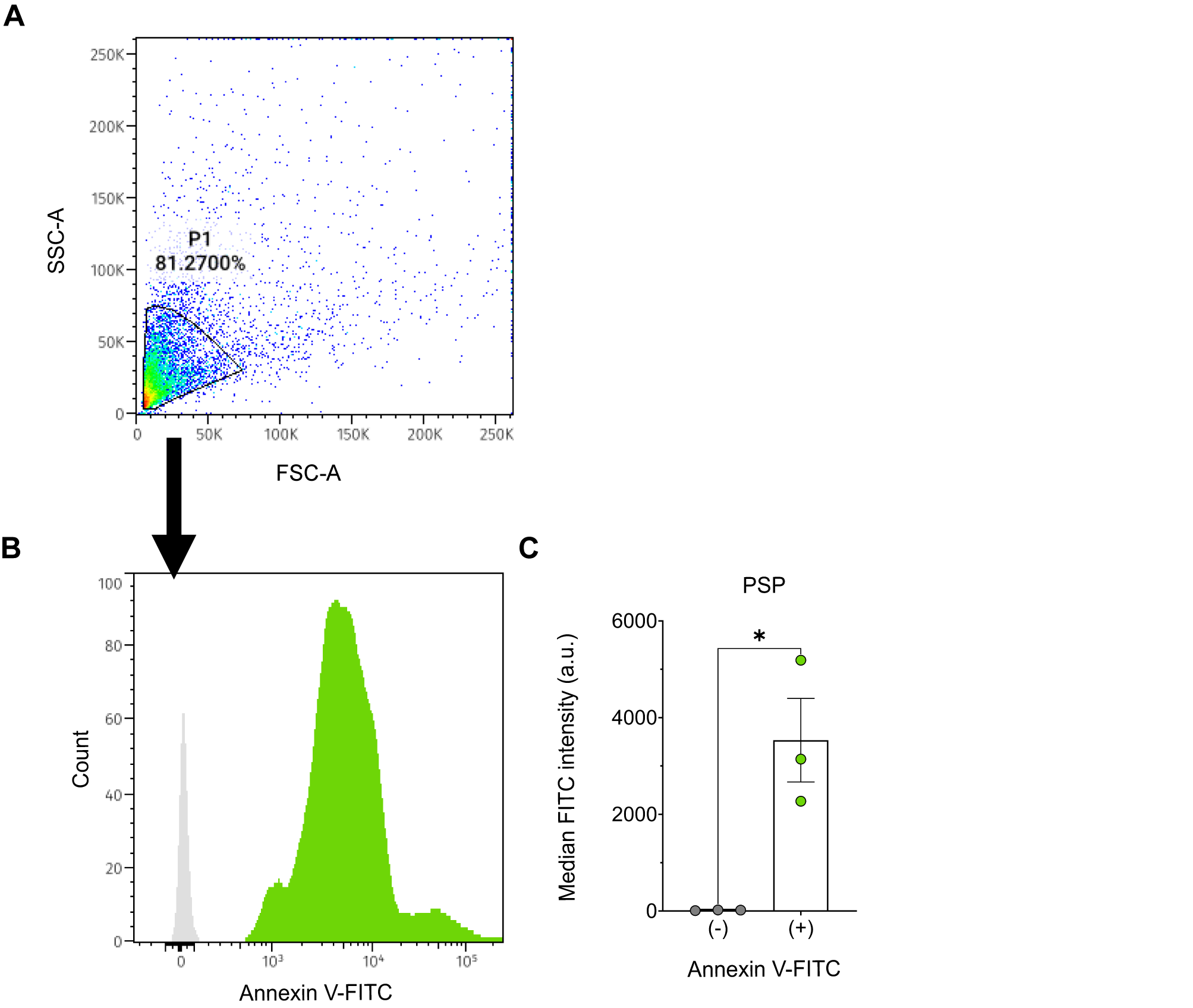

**Fig. S2.** **PS validation using Annexin V-FITC assay**. **A-B**, Representative FSC-A vs. SSC-A gating (**A**) and histogram (**B**) of PSP with or without Annexin V-FITC. Grey represents PSP only (background), while lime green shows PSP + Annexin V-FITC. A clear rightward shift (lime green) relative to background (grey) is observed indicating the positive detection of PS. **C**, Median FITC intensity of PSP with or without Annexin V-FITC. Data represent means ± SEM (n = 3 independent particle preparations).

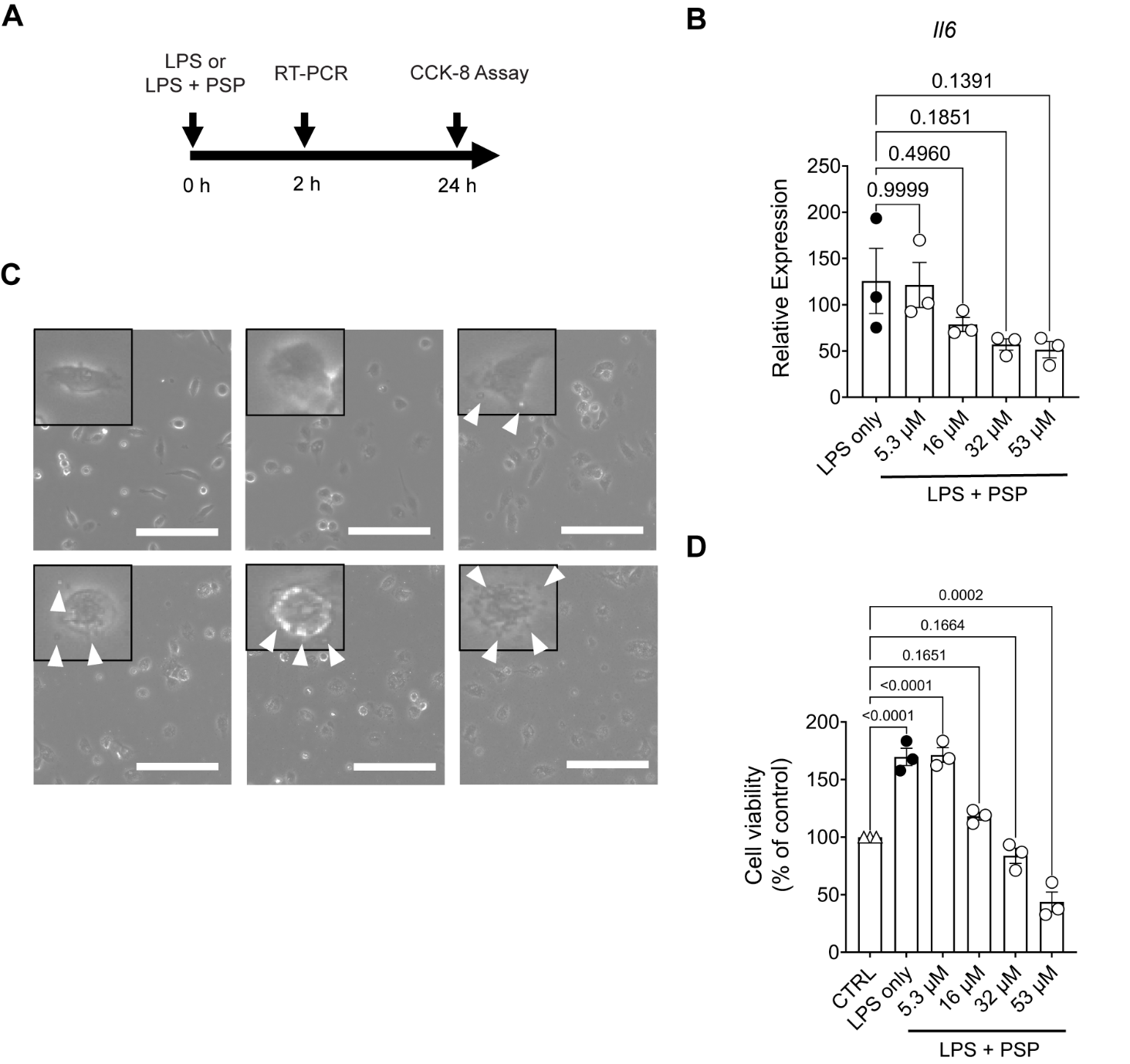

**Fig. S3. PSP downregulates *Il6* in a dose-dependent manner.** **A,** Evaluation scheme of PSP for mRNA expression and cell viability assay. **B,** Treatment with PSP at various concentrations. Data represent means ± SEM (n = 3 biologically independent animals). **C,** Brightfield images of cells at various treatments. Inset corresponds to magnified image of a single cell, while arrowheads point to visible particles. Scale bar = 100 µm. **D**, Cell viability of macrophages at various PSP concentrations. PSPs used were 500 nm. CTRL: untreated samples. Data represent means ± SEM (n = 3 biologically independent animals). All graphs were analyzed using one-way ANOVA.

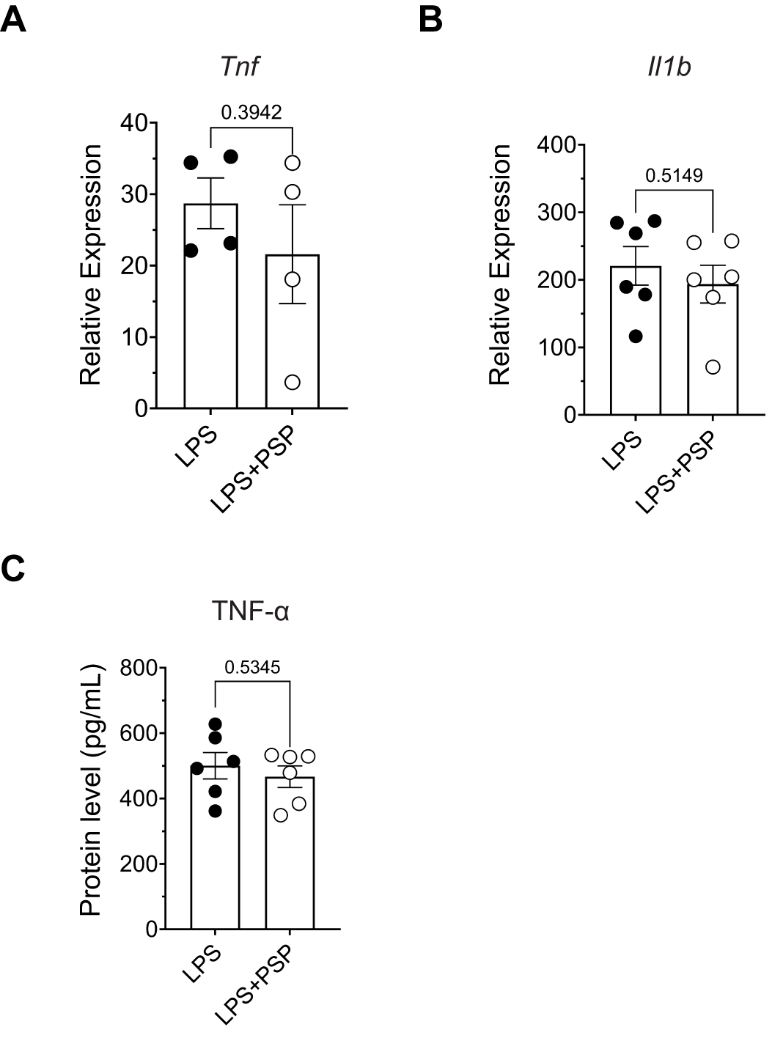

**Fig. S4. Evaluation of PSP effect on other pro-inflammatory cytokines.** **A-B,** Gene expression level of *Tnf* (**A**) and *Il1b* (**B**). **C,** TNF-⍺ level. Data represent means ± SEM (n = 4 to 6 biologically independent animals). All graphs were analyzed with two-tailed unpaired t-test.

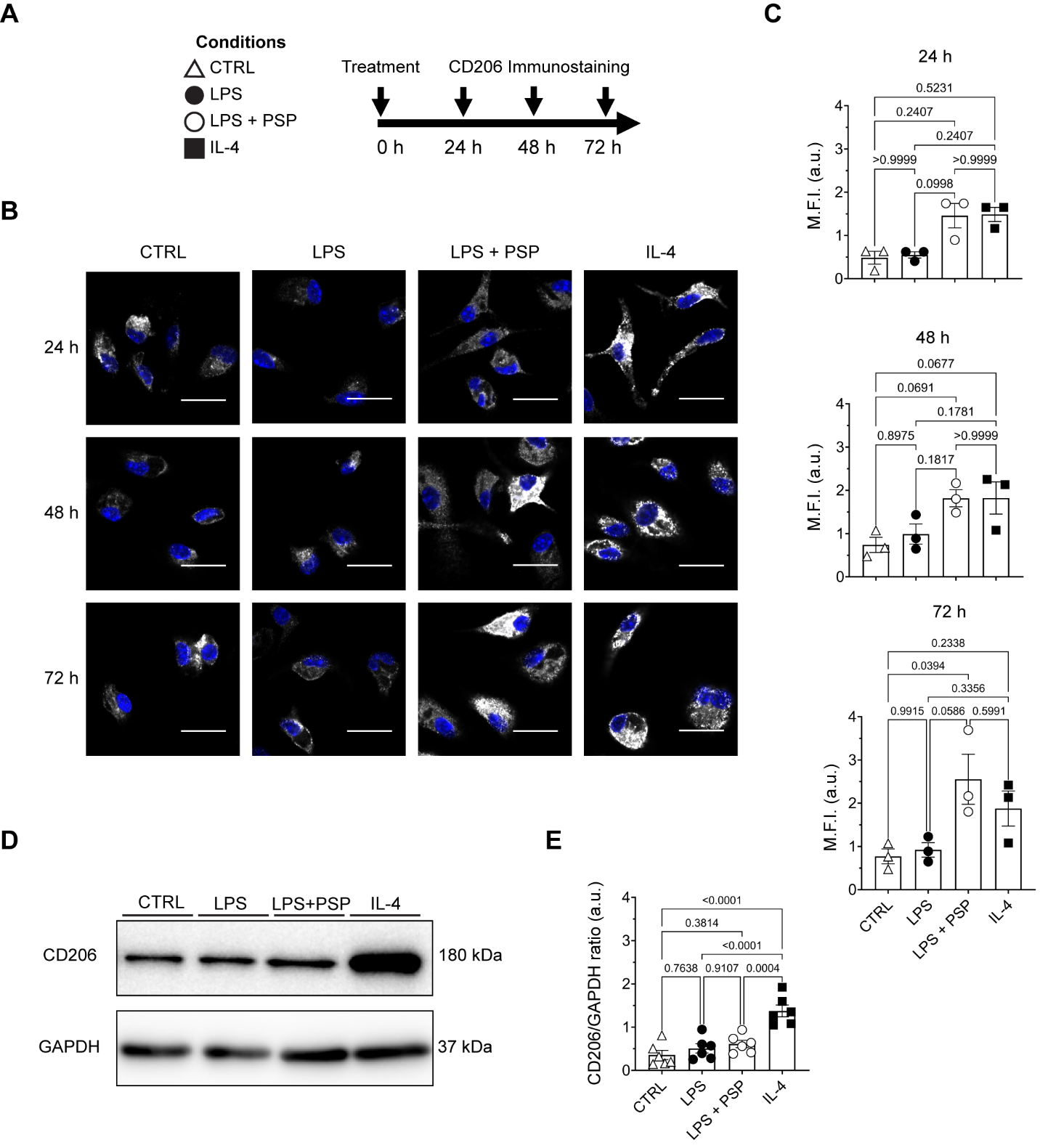

**Fig. S5. Evaluation of CD206 expression following PSP treatment. A**, Evaluation scheme of PSP for M2 macrophage marker. CTRL: untreated samples. **B**, Immunostaining images of different treatment conditions at different time points. Scale bar = 20 µm. **C,** Quantification of CD206 Mean Fluorescence Intensity (M.F.I.). Data represent means ± SEM (n = 3 biologically independent animals). **D**-**E,** Western blot of total protein extracts. Data represent means ± SEM (n = 6 biologically independent animals). All graphs were analyzed using one-way ANOVA, except in 24 h (top panel, **C**), where a Kruskal-Wallis test was performed.

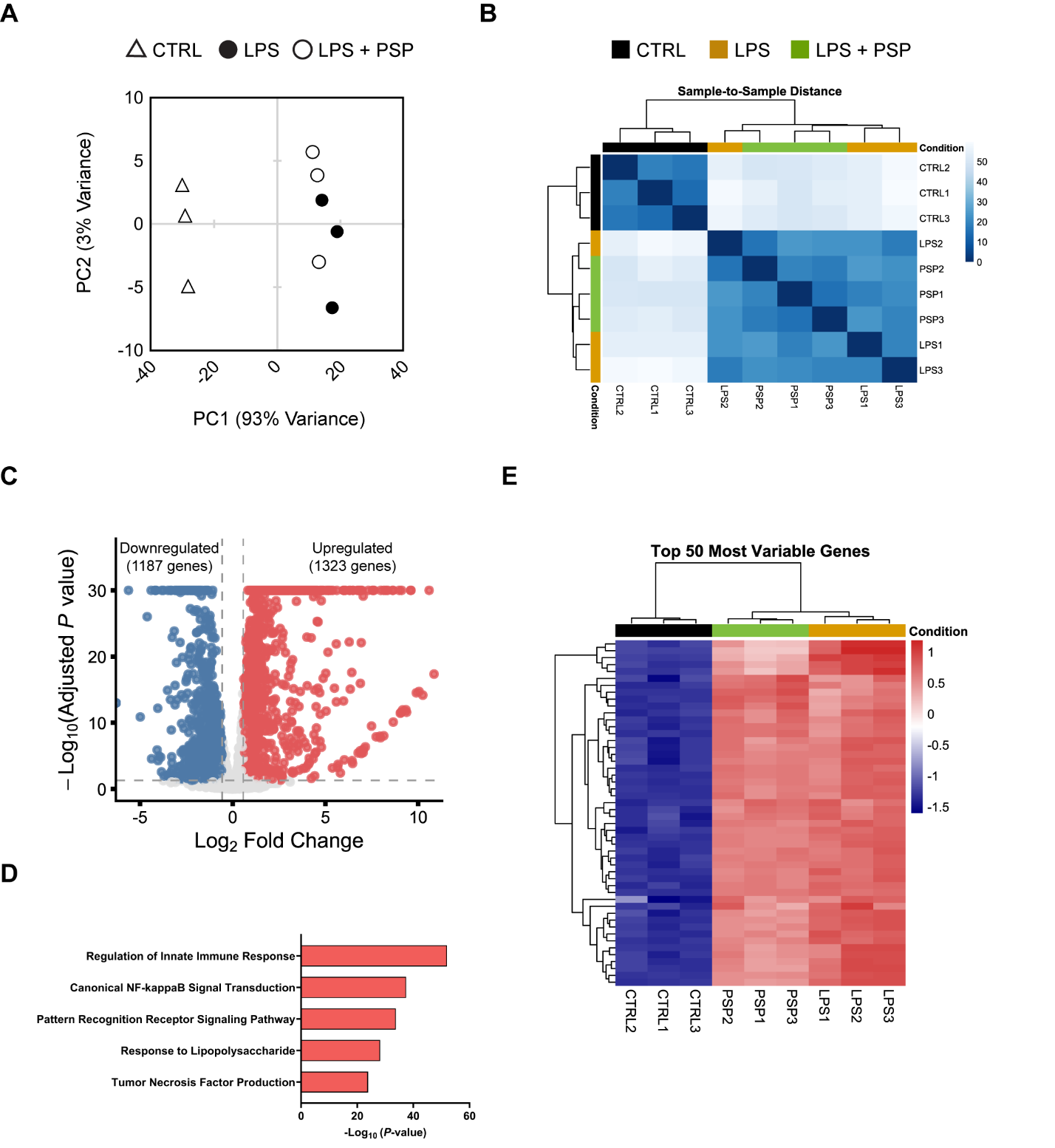

**Fig. S6**. **Transcriptomic analysis of untreated and treated groups.** **A**, Principal Component Analysis (PCA). CTRL: untreated samples. **B,** Sample-to-sample distance heatmap. **C,** Volcano plot showing the differentially expressed genes (DEGs) from LPS vs. CTRL. **D**, Enriched pathways induced by upregulated DEGs (1323 genes)**. E**, Hierarchical clustering heatmap of the top 50 Most Variable Genes.
