## Supplementary Table for "Immunomodulatory mechanisms of submicron phosphatidylserine-exposing polymeric particles (PSPs)"

**Table S1.** RT-PCR primers

| *Actb* | Forward | 5' - GATCAGCAAGCAGGAGTACGA - 3' |
| --- | --- | --- |
|  | Reverse | 5' - AAAACGCAGCTCAGTAACAGTC - 3' |
| *Il1b* | Forward | 5' - TGCCACCTTTTGACAGTGATG - 3' |
|  | Reverse | 5' - TGTGCTGCTGCGAGATTTGA - 3' |
| *Il6* | Forward | 5' -CACTTCACAAGTCGGAGGCT - 3' |
|  | Reverse | 5' -CTGCAAGTGCATCATCGTTGT - 3' |
| *Tnf* | Forward | 5' - AGCCGATGGGTTGTACCTTG - 3' |
|  | Reverse | 5' - ATAGCAAATCGGCTGACGGT - 3' |
